# Rethinking data treatment: The sucrose preference threshold for anhedonia in stress-induced rat models of depression

**DOI:** 10.1101/2023.03.31.535101

**Authors:** Jenny P. Berrio, Otto Kalliokoski

## Abstract

Exposing rats to repeated unpredictable stressors is a popular method for modelling depression. The sucrose preference test is used to assess the validity of this method, as it measures a rat’s preference for a sweet solution as an indicator of its ability to experience pleasure. Typically, if stressed rats show a lower preference compared to unstressed rats, it is concluded they are experiencing stress-induced anhedonia. While conducting a systematic review, we identified 18 studies that used thresholds to define anhedonia and to distinguish “susceptible” from “resilient” individuals. Based on their definitions, researchers either excluded “resilient” animals from further analyses or treated them as a separate cohort. We performed a descriptive analysis to understand the rationale behind these criteria, and found that the methods used for characterizing the stressed rats were largely unsupported. Many authors failed to justify their choices or relied exclusively on referencing previous studies. When tracing back the method to its origins, we converged on a pioneering article that, although employed as a universal evidence-based justification, cannot be regarded as such. What is more, through a simulation study, we provided evidence that removing or splitting data, based on an arbitrary threshold, introduces statistical bias by overestimating the effect of stress. Caution must be exercised when implementing a predefined cut-off for anhedonia. Researchers should be aware of potential biases introduced by their data treatment strategies and strive for transparent reporting of methodological decisions.

## Background

Testing for sweet preference is a popular method for assessing anhedonia in rodent models of depression. Rodents naturally like sweet things, and when presented with two bottles, one containing a sweetened solution and the other water, they more often than not drink from the former. The practice of presenting a rat with two water bottles where one has been sweetened, and measuring the consumption of each, is commonly known as the sucrose preference test (Figure 1). When given free access to the bottles, on average, 80% of a healthy rat’s total liquid intake will come from the sweetened solution^1–3^. When studies find that the preference changes after, for example, chronic stress, the change is thought to reflect a reduced responsiveness to rewards. The once pleasurable experience of consuming something sweet is no longer as rewarding. This, by definition, is anhedonia, a core symptom of depression. In stress-induced models of depression, such as the chronic unpredictable stress model^4,5^ the sweet preference of stressed animals is often measured. When stressed subjects consume significantly less sweet solution than do unstressed conspecifics, it is suggested that they have comparatively less capacity to experience enjoyment – they are experiencing stress-induced anhedonia^4^. However, this relatively simple idea becomes more complex when we consider the details. For example, by how much would a rat have to reduce its intake of sweetened solution for us to consider it to be experiencing anhedonia?

**Figure 1.**
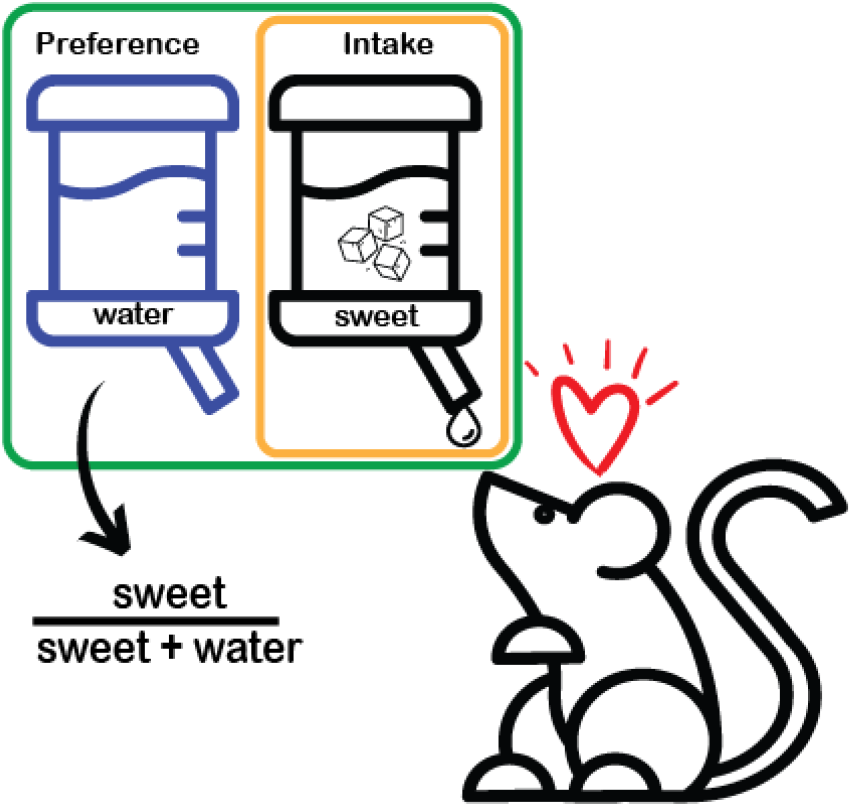
Simplified illustration of the sucrose preference test. In a typical test, two bottles are presented to the rat. One contains plain water while the other a sweetened solution, commonly a sucrose solution (1-2% w/v). After a period that allows for consumption (frequently 1 h), the intake or the preference is recorded and compared between groups of interest. Intake is simply the amount of sweetened solution consumed, while preference (formula in the illustration) is the proportion of the consumed liquid that was made up by the sweetened solution. Copyright attribution can be found in the supplementary material.

### Defining Anhedonia

We recently reviewed 159 studies (where 132 presented data that could be analyzed) of laboratory rats that were experimentally stressed for two weeks or more, and that were assessed for anhedonia using the sucrose preference test. We focused on studies that used no fasting, or only short periods of food and/or water deprivation, before the test. These studies constituted only 18.3% of the total studies we found during our search that used the model^6^. In 18 of the 159 studies, the authors set a criterion for defining anhedonia. That is, they set a specific sucrose consumption threshold for determining whether a stressed animal was experiencing anhedonia or not. This is critical since not all rats reduce their sweet intake in response to stress. In some instances, these stressed animals – considered non-anhedonic – were excluded from further analysis, focusing the study only on animals considered “susceptible” to the stress protocol^7–14^. In other studies, the non-anhedonic animals formed a new cohort and were considered “resilient” to the effects of stress on reward sensitivity^15–24^. The concept of susceptibility and resilience to stress is not new. In both humans and animals, the mechanisms leading to a differential response to stress are of great interest and an important focus of research^25–27^. How this distinction is made, however, has implications for the conclusions drawn from any analysis – whether behavioral or physiological – comparing the resulting populations. Yet, during our review process, we could not find one common, agreed-upon, criterion for defining anhedonia in the sucrose preference test; instead we found many. In the interest of knowing how these different criteria were chosen, we performed a descriptive analysis of these 18 studies.

## Methods

We searched for how susceptibility to anhedonia was defined in terms of the sucrose preference test, as well as any methodological or biological justification for it, in all sections of the selected study reports. In cases where this justification was provided in the form of references to previous publications, the cited material was back-traced similarly to what has been described by Standvoss and colleagues^28^. Briefly, the referenced source was retrieved and if it, in turn, did not provide a justification but further references, the process was repeated. Sources were retrieved until a justification was found or a dead end in the citation chain was reached. Dead ends occurred when no justification was found, neither in the text nor in referenced form. The sources accessed to reach a justification, or a dead end, were recorded, along with their type (original study/review).

## Results

The results of this analysis confirmed our first impressions (Table 1). How susceptible animals were defined varied greatly among the screened studies. Three general methods were identified for separating rats into the two sub-populations. Four studies compared the performance of stressed rats to that of control animals, not only in the sucrose preference test^15^, but also in other behavioral tests such as the open field test and the forced swim test^7,18,21^. Sensitivity to stress was defined as an animal exhibiting abnormal behavior in two or more of the tests. Only one study presented a clear definition of what was considered abnormal^7^. Three studies separated their animals based on a within-subject decrease in sucrose consumption^16,17^ or preference (the proportion of the total liquid intake that was made up of the sucrose solution)^19^. How much rats had to change their sweet intake from their initial values to be considered stress-susceptible varied from 25% to 50%. The criteria for resilience was 10% or less. Most studies (11) established a cut-off based on sucrose preference (Table 1). Most studies (11) established a cut-off based on the sucrose preference (Table 1). Rats with preference below a certain threshold were classified as susceptible, whereas those with values above were classified as resilient. This threshold varied from 55% to 75%, but was most frequently set at 65% (7 studies). Despite analyzing different studies, our findings align with a recent review of the most influential (most cited) studies using chronic unpredictable stress in rats^26^. This review also found different means of stratifying animals into groups of susceptible and resilient individuals. This finding suggests that the observed heterogeneity in classification we identified in our study is widespread in the field, rather than being specific to the subset of studies we analyzed.

**Table 1.**
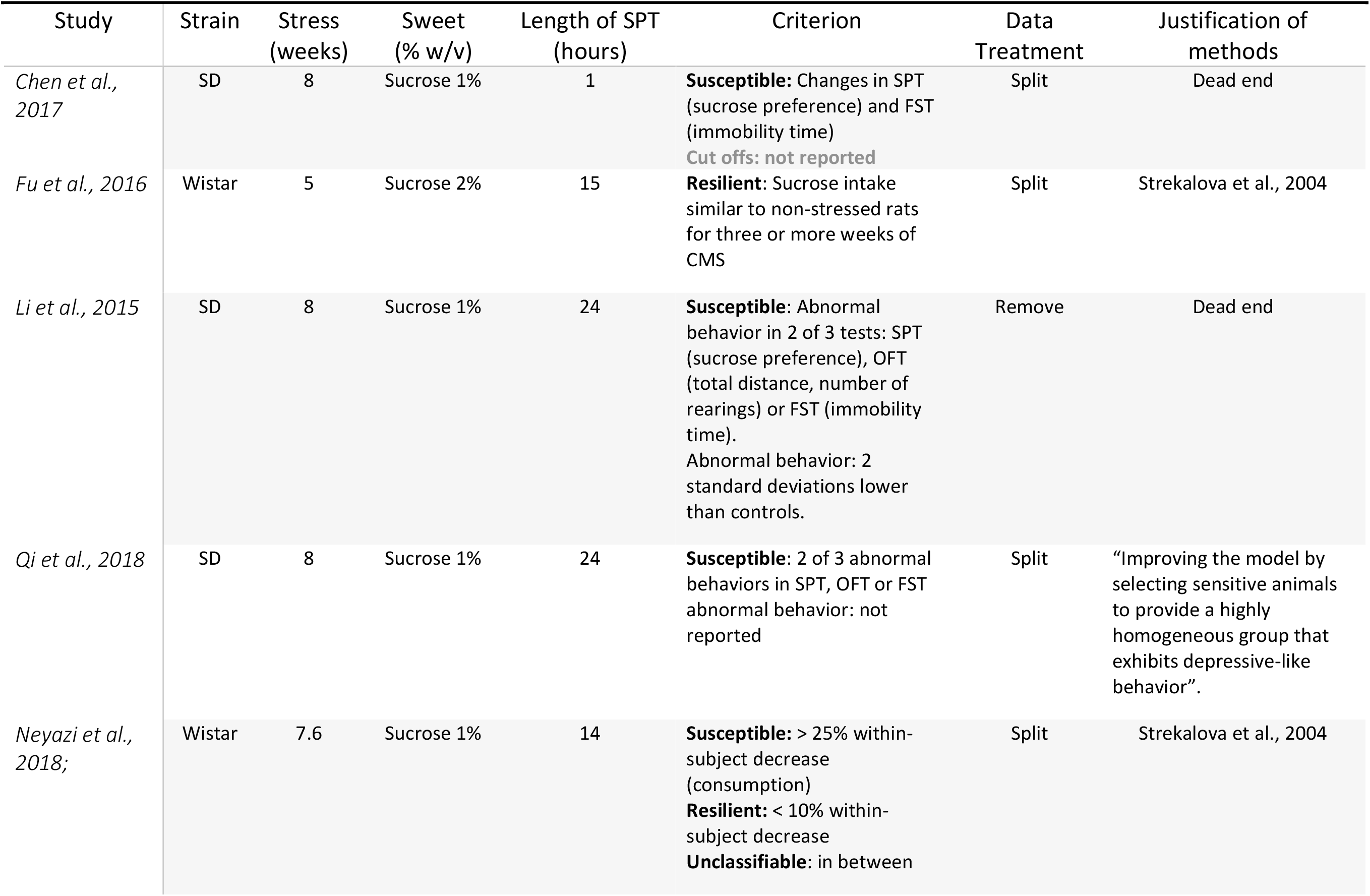
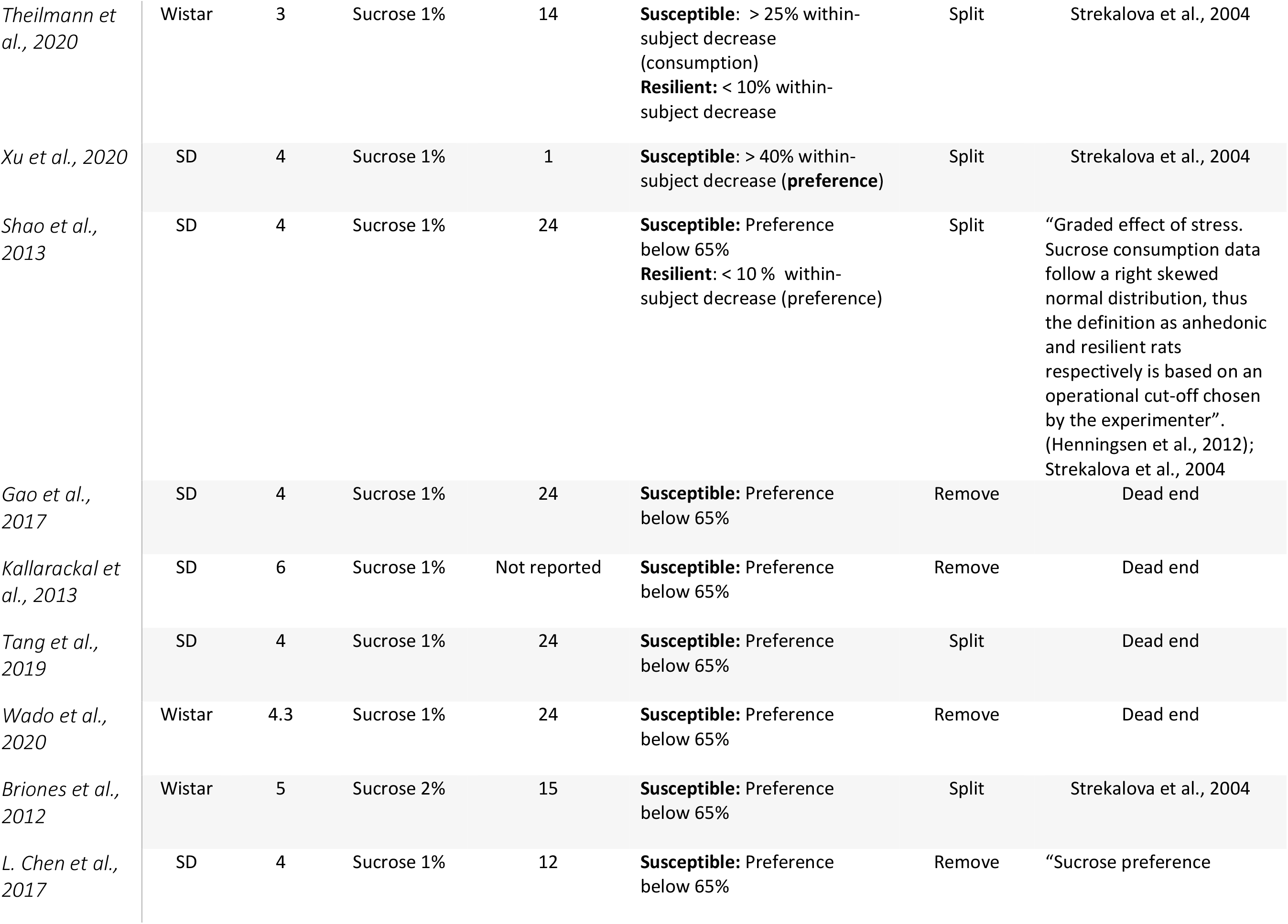
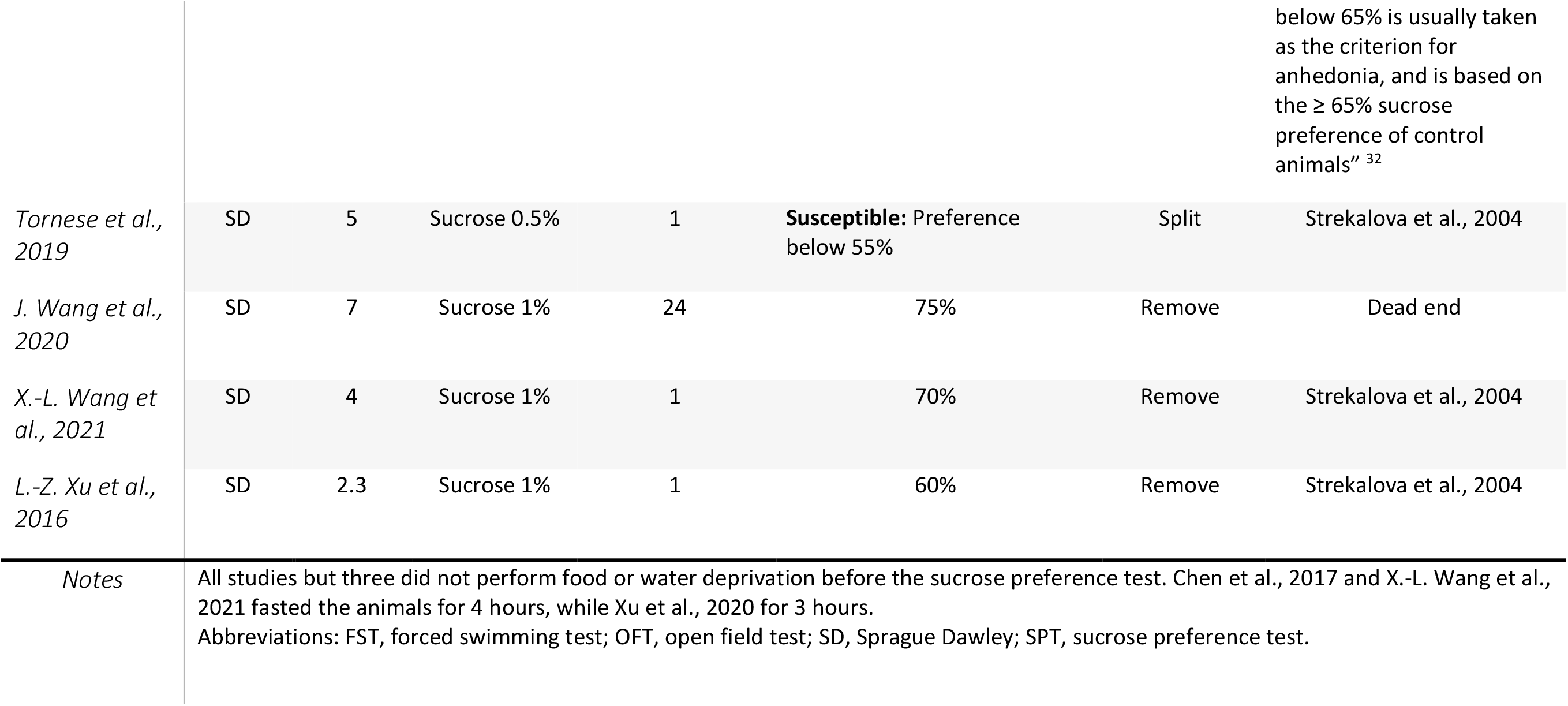
Descriptive summary of back-tracing experiment.

While criteria were varied, justifications were scarce, and in most cases authors relied on citing previous research without further explaining the rationale behind their method. In 38.9% of the studies (7/18), no citations or justifications were provided. In the remaining studies, we often found cross-referencing among papers employing different methods of classification. Some studies cited reports using similar strategies, but then employed different cut-off values. In other instances, studies employing a preference threshold were cited when, instead, a change in consumption from baseline was opted for. No explanations for these disparities were found. We were able to, ultimately, identify three different justifications in our investigation. Qi and colleagues (2008)^18^ motivated their combined use of three tests, noting that they would collect a highly homogeneous group that expressed similar depression-like behaviors in this way. Henningsen and collaborators^34^ provided a methodological explanation to their choice of a minimum change in consumption from baseline: “the sucrose consumption data follow a right skewed normal distribution, thus the definition as anhedonic and resilient rats respectively is based on an operational cut-off chosen by the experimenter (30% within-subject decrease)”. Since no other study using this methodology presented additional explanations, we can only speculate whether such an operational choice was also used by others. Hurley and collaborators^32^ argued that 65% is often chosen because unstressed controls have preferences above this value. This justification mirrors one provided by a paper written by Strekalova and collaborators (2004)^30^ which was frequently reached in the chain of citations justifying the criteria for susceptibility and resilience (Figure 2).

**Figure 2.**
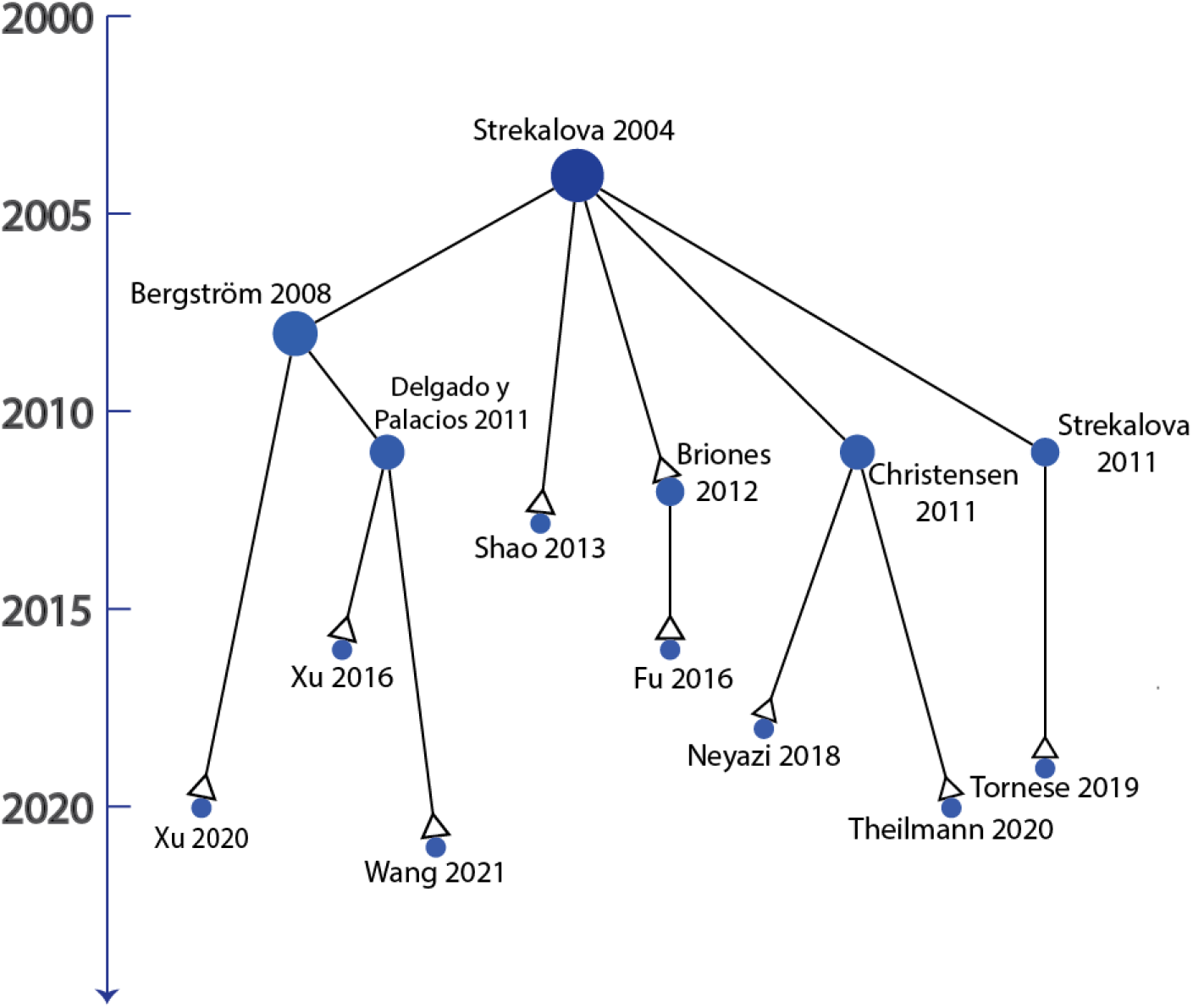
Simplified back-tracing timeline. Nine of the 18 studies directly or indirectly cited Strekalova *et al*., 2004. These studies are identified with a white arrow in the graph. The timeline in years is represented on the Y-axis.

This central paper is the first in a series of studies^35^ that attempted to distinguish sub-populations of mice subjected to stress, focusing specifically on anhedonia. Anhedonia was determined using the sucrose preference test, with a cut-off for the preference level set at 65%. The authors explained that this choice was based on two facts. One: none of their unstressed mice had a preference below 65% after four weeks of stress. Two: Rats and mice with a preference below this value had shown other features of anhedonia and depression in previous studies. However, when the supporting references were followed, we found no direct (published) evidence supporting this later claim. All but one^36^ of these citations are studies in rats. Only two of the cited studies assessed the effects of chronic unpredictable stress using the sucrose preference test^36,37^, of which only one reported the outcome in the form of preference^36^. This study made no distinction between different responders to stress. Moreover, the study^38^ cited as evidence that rodents with a preference below 65% also show an increased intracranial self-stimulation threshold did not actually report on having performed a sucrose preference test. Nor did studies conducted by the same research group up to 1995^39–41^. So, while often cited as support, the pioneering approach by Strekalova and collaborators cannot be considered an evidence-based method for separating stress-sensitive and stress-susceptible mice. To this we will add that mice and rats are different species; species that respond differently in behavioral tests like the sucrose preference test.

## Discussion

We do not aim to undermine the valuable research that has been done in characterizing susceptibility and resilience to stress. However, we would like to caution against implementing a specific cut-off in the sucrose preference test to define anhedonia based solely on previous literature. Particularly since this practice is not restricted to the small subset of studies we found through our review, but it is prevalent in the field. The chronic unpredictable stress model of depression, while popular, has a disadvantage in that the smallest changes in the methodology can cause great variability in the observed sweet consumption^5,42^. Differences in stress protocols and flexibility in testing methods, particularly in the case of sucrose preference testing, has led to high heterogeneity in the phenotype of the model^6,43^. It is not uncommon to find two studies with differing durations in fasting before the test, length of the test itself, or the concentration and type of the sweet solution employed. In our review, we found the sucrose concentration to range from 0.2% to 30% (w/v) and the length of test from 15 minutes to 8 days^6^. Different sucrose concentrations and test durations were also employed in the studies that are the center of the present paper (Table 1). Rodents’ preference for sweet things varies in relation to the type of sweetener being used and its concentration^3,44,45^. Indeed, some rodents are not particularly enticed by sweetness to begin with^2^. In our data, we also found high variation in the preference for sucrose of control groups (Supplementary figure 1). Since individual variation is unavoidable, and conditions across laboratories are different even when a similar model and test are employed, a universal cut-off seems counterintuitive.

More importantly, while theory assumes that a decrease in sweet consumption reflects a diminished ability to experience pleasure (anhedonia), this has never been proven. A study that assessed sucrose consumption and self-stimulation thresholds in the same groups of animals failed to show that a decreased sucrose intake following chronic stress is accompanied by an increased intracranial self-stimulation^46^. In our review^6^, we found evidence of non-hedonic factors driving the consumption of sweet solutions. What is more, we observed that methodological strategies, meant to reduce bias when conducting the test (for example diminishing the effect of side preference on the test results), were often not implemented or not reported on. With this in mind, can we be certain that a changed behavior in the sucrose preference test actually reflects a change in the hedonic state of an animal and is not simply the result of one or more unrelated factors?

It should be noted that there are different subtypes of anhedonia – facets linked to liking, wanting and reward-related decision making; each governed by specific, although interlinked, neurobiological processes^47^. The sucrose preference test is ostensibly a measure of liking (consummatory anhedonia) rather than wanting (motivational anhedonia) and it would be incorrect to assume that a deficit in a subtype of anhedonia must reflect a general deficit. What is more, caution must be taken when translating the results of a test measuring a subtype of anhedonia in animals to deficits in humans, in whom, the wanting component is mostly researched^48^ and the processes linking liking and wanting (decisional anhedonia)^47^ are much more complex. If used side by side with other measures of response to rewards (*e*.*g*., intracranial self-stimulation) we can gain a better understanding of what the sucrose preference test measures and its limitations. Until then, relying on a particular threshold of consumption/preference as a direct measure of general anhedonia risks reaching misleading conclusions.

Another important point we would like to make concerns quality of reporting. Scientists characterizing different stress-induced behavioral phenotypes would do well to improve their methodological reporting. We found that authors had a tendency to rely on citations as the sole means of contextualizing or justifying the criteria used to define anhedonia. This happened even when the criteria in the cited material differed from their own. We are not alone in our findings. A recent study by Standvoss and colleagues (2022) determined that between 19% and 26% of over ten thousand citations found in the methods sections of 815 papers in the fields of neuroscience, biology and psychiatry aimed at providing context or explaining why a particular method was chosen. While this shortcutting is not a bad practice *per se*, it does come at a cost for the reader and may affect reproducibility if not done effectively^28^. Tracing chains of citations is a time-consuming task. Even more so when multiple citations are provided. Consider for a moment diligently tracing, accessing and reading all citations, only to find an unexplained discrepancy between the citing paper and the cited material in crucial aspects of the method. Failing to address the rationale behind methodological mismatches, not only wastes fellow researchers’ precious time, but also potentially denies them key information. This lack of transparency can hinder scientific progress and lead to inconsistencies in research findings.

### Data treatment strategies and their impact

The way we analyze data obtained in the sucrose preference test, once a cut-off for anhedonia has been selected, can introduce statistical bias. Eight of the 18 papers we have chosen to focus on removed animals that were not considered anhedonic from further analysis (Table 1). This practice not only discards valuable information that might help in understanding the processes leading to unexpected results after stress (no change in the intake of sweet solution, or an increased consumption), but also exaggerates the effects of stress. The other ten studies separated the stressed animals into two subgroups (susceptible, resilient), which were later compared to control animals that had not been exposed to stress (Table 1). While this practice retains key information, it still increases the risk of false discoveries. In order to understand the effect of these analytical choices, we simulated the sucrose intake for cohorts of unstressed rats. When comparing these groups to one another, any statistically significant differences we might find will have arisen only by chance, as the effect of stress was not incorporated in the simulation.

## Methods

We simulated the results from a sucrose preference test for cohorts of 12 animals, where each animal’s sucrose intake was based on values reported in the literature. We then compared two of these cohorts to one another. Repeating this process over and over would give us an approximation of the false positive rate (how often a statistically “significant” difference was found between the two groups with respect to their sucrose preference) for experiments where we did not differentiate individuals according to susceptibility to anhedonia. We then looked at the two analytical choices we found in our literature searches that introduced a cut-off, subdividing cohorts into susceptible and resilient individuals (Figure 3). In the first case, one of the cohorts was modified by simply removing the “resilient” rats whose sucrose preference exceeded 65% (recall that these are unstressed control rats, and consequently there is nothing to which they should be resilient). The unmodified cohort was then compared to the modified cohort using a simple t-test. In the second case, one of the cohorts was split into two sub-cohorts. Rats with a preferences above 65% formed a “resilient” cohort, whereas the others became a “susceptible” cohort. The unmodified cohort and the two modified sub-cohorts were compared using a one-way ANOVA. We report values based on the results of 1,000,000 simulated experiments. Our simulations used cohorts of 12 animals and a cut-off set at 65%, since our review of the literature pointed to this being a typical design for a study using the sucrose preference test. The code for the simulations was created using RStudio 2022.07.2.576^49^ with R version 4.2.1^50^ and it is accessible through (https://osf.io/egk8s/). The simulated preferences (Figure 4) were based on values for sucrose and water consumptions of unstressed rats that were extracted from the dataset of our systematic review (Supplementary material). We chose to use data from tests using sucrose solution (1-2% w/v) in 1-hour tests as this is the most typical mode of testing used among researchers^5^. A sucrose intake of 12.22 ± 7.49 (mean ± SD), water intake of 2.66 ± 1.63, and leakage (mL) per bottle of 1 to 3 droplets was assumed for the model. We made the code accessible to any user through a user-friendly and free Shiny app (http://jenb.shinyapps.io/App-1). The app offers users the flexibility to explore the effects of the two outlined data treatment strategies using either their own sweet consumption data or the data we have retrieved through published sources. The app was named the “Sweet preference test simulator” to emphasize that it can be used with various types of sweeteners including, but not limited to, sucrose. For more detailed information on the app and a complete list of the packages used, see the supplementary material.

**Figure 3.**
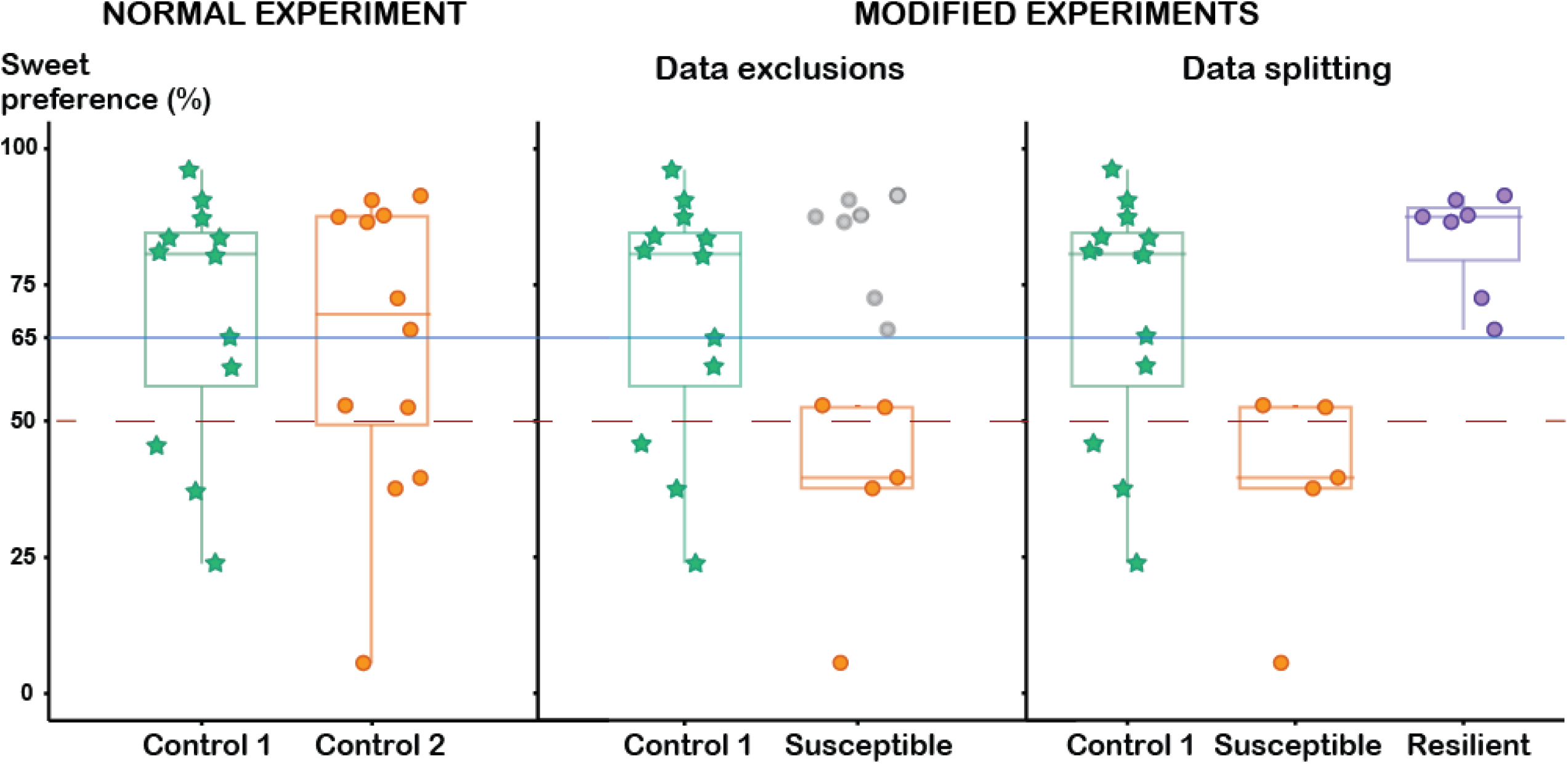
Graphical representation of the three types of experiments we simulated. *Normal experiment:* Two cohorts of twelve unstressed rats each (referred to as control 1 and control 2) are compared on their simulated preference using a simple t-test. Any significant differences that may arise are only a product of chance, since the effects of stress are not simulated. *Data exclusions:* the cohort 2 is modified by removing rats with preferences above 65% (blue line - grey data points), then the modified cohort (referred here as “susceptible”) is compared to the unmodified control 1 with a simple t-test. *Data splitting:* the cohort 2 is split into two: Rats with preferences above 65% form a “resilient” cohort, whereas those below 65% are labelled “susceptible.” The resulting three cohorts are compared using a one-way ANOVA. Values above 50% indicate a preference for the sweetened solution over water (dashed red line).

**Figure 4.**
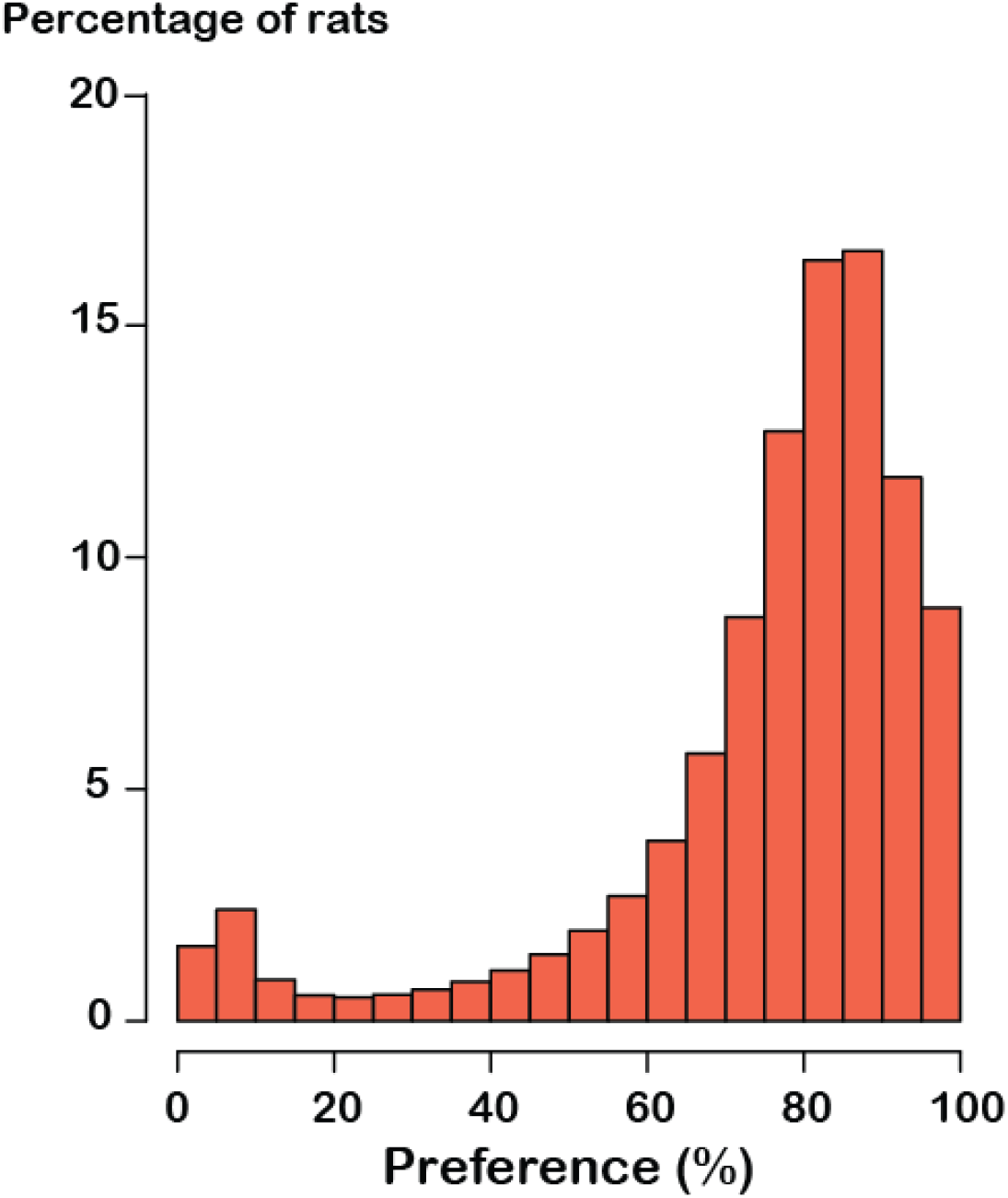
Histogram of the preference values for simulated rats. We have simulated the results for one million unstressed control rats.

## Results

In our stochastic simulation, 3.7% of the normal experiments found a difference between the un-modified cohorts when there was none (Figure 5A). With a cut-off as low as 65%, a number of the experiments in which one cohort was modified (59% of simulated experiments) could not be analyzed statistically since fewer than three rats were categorized as “susceptible.” In the experiments where we could test the two data treatment strategies we saw a greatly increased rate of false positive experiments. In half (49%) of the experiments that excluded “resilient” rats, the cohorts were found to be significantly different (Figure 5B). In 71% of the experiments in which one cohort was split into two, the unmodified cohort and the “susceptible” cohort were found to be significantly from one another (Figure 5C). Moreover, the proportion of viable experiments – experiments where a statistical comparison was possible (*i*.*e*., all cohorts consisted of more than three animals) – that found a false positive increased with greater sample sizes (Figure 6 A, B) and with lowered preference cut-offs (Figure 6 C, D).

**Figure 5.**
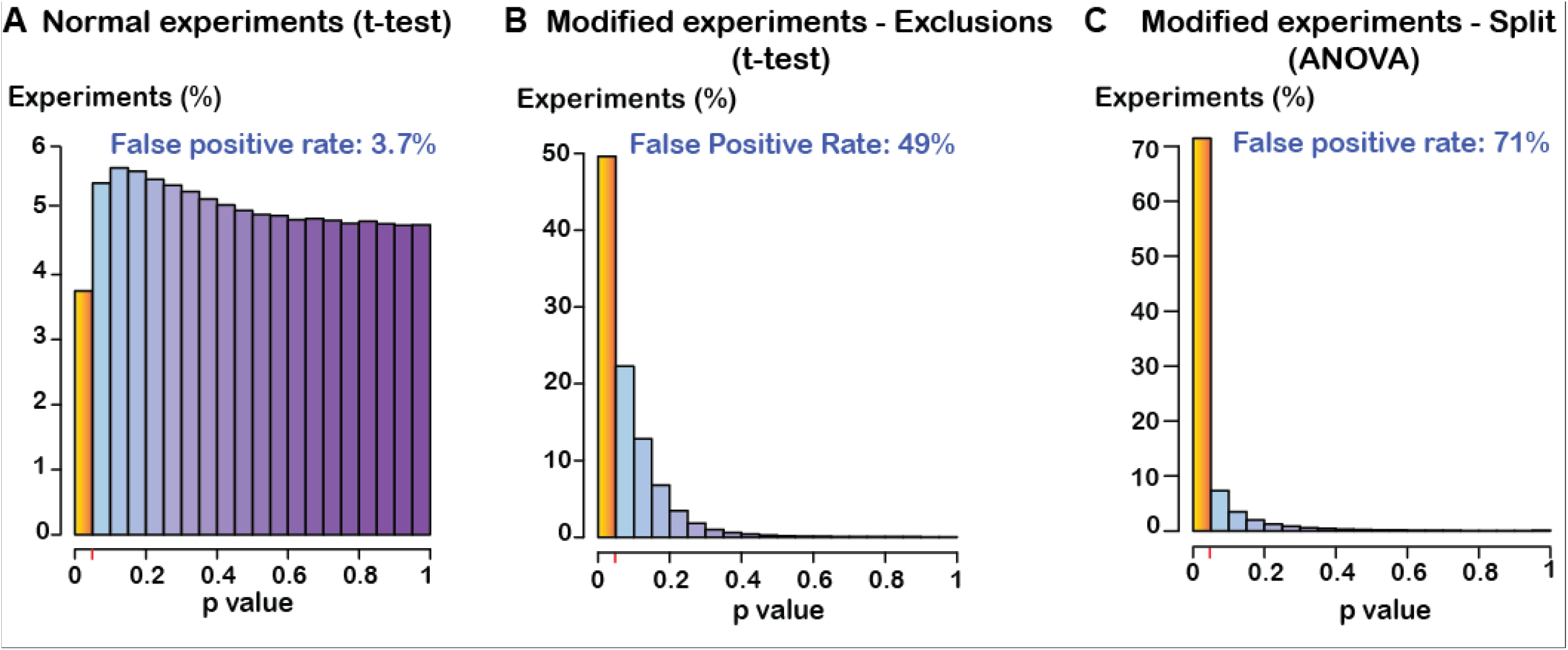
False positive rates in three types of simulated experiments. A. Experiments in which two unmodified cohorts of rats were compared to each other using a t-test. The false positive rate is lower than the expected 0.05. This is due to the non-normal distribution and small sample sizes. When larger sample sizes were used, this value matched the 5% alpha level (Supplementary figure 2). **B**. Experiments in which one of the cohorts was modified by removing “resilient” individuals with a sucrose preference above 65%. Each modified cohort was compared to an unmodified cohort using the t-test. **C**. Modified experiments in which one of the cohorts was split into two. Rats with a sucrose preference above 65% formed a “resilient” sub-cohort, whereas the others formed a “sensitive” sub-cohort. The sub-cohorts were compared to an unmodified cohort in a one-way ANOVA. The false positive rate is calculated for experiments where statistical comparisons were possible. In 59% of the experiments these comparisons were not possible after the cohorts had been modified, as less than three rats remained in the sensitive sub-cohort. Red markers on the X-axis/yellow bars represent a p value of 0.05.

**Figure 6.**
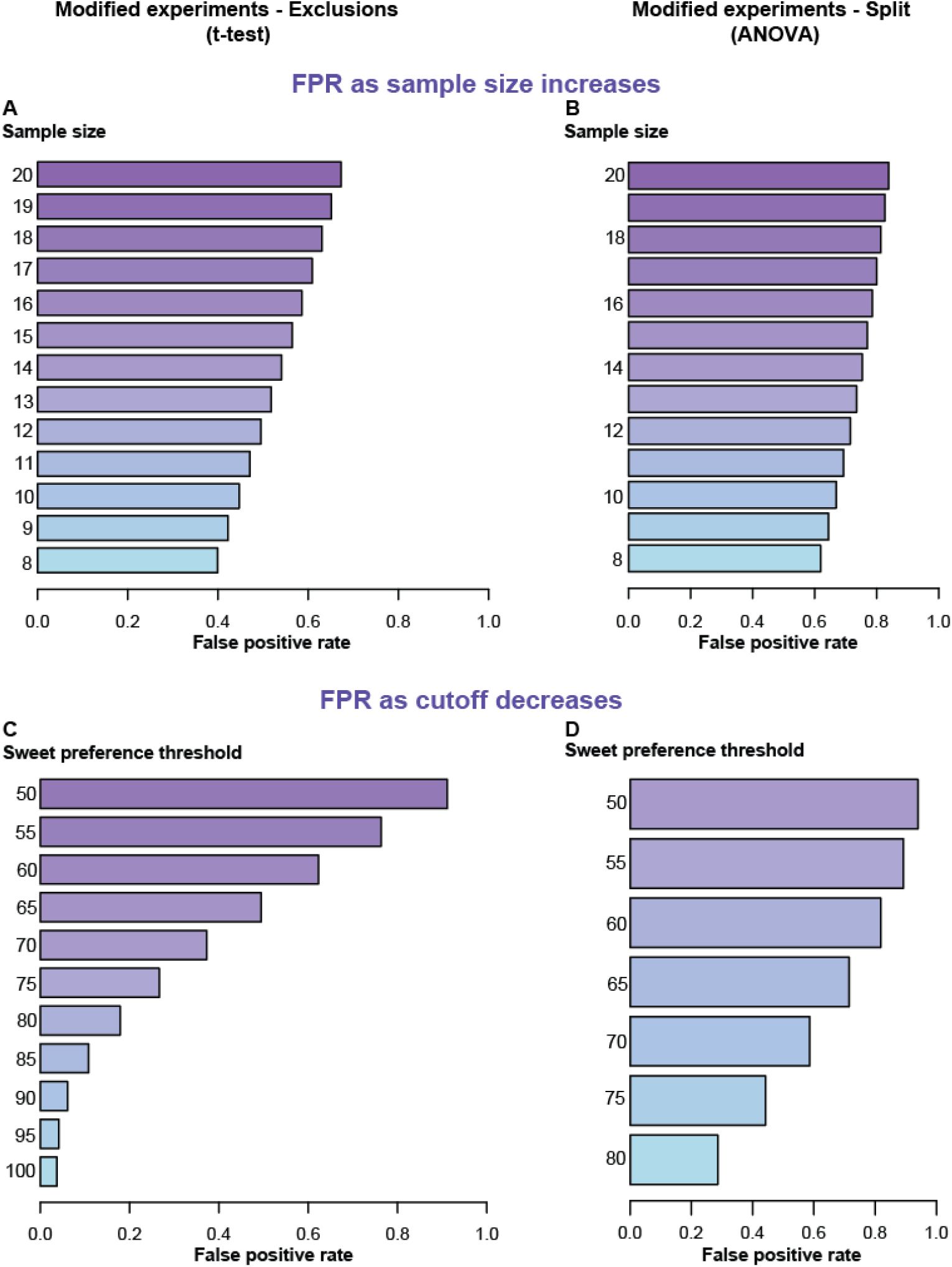
False positive rates with greater sample size and different preference cut-offs. Modified experiments in which “resilient” rats were removed (A, C). Modified experiments in which one of the cohorts was split into two (“susceptible”, “resilient”), and an unmodified cohort was compared to the two sub-cohorts. The false positive rates are of the *post hoc* comparisons between the unmodified cohort and the “susceptible” sub cohort (B, D).

## Discussion

Our simulations demonstrate that, not only we are relying on the yet-to-be proven assumption that the sucrose preference test is a reliable measure of general anhedonia, but that frequently-used data treatment strategies further increase the risk of reaching misleading conclusions. Even in the absence of the effect of stress, these strategies would frequently find significant differences between groups of individuals sampled from the same population. In the real-world context, these differences would have been interpreted as an effect of chronic stress. In cases where stress does have an effect on sucrose consumption, practices like these may confound the results of the experiment since they overestimate this effect. In other words, researchers may unintentionally end up with a result that is not as reliable or accurate as it could have been. Unexpectedly, we found the rate of false positive experiments to increase, roughly in a linear fashion, with increasing sample size. This may seem counterintuitive since larger sample sizes in most cases yield more robust findings. With large enough samples, researchers may feel confident that removing a few data points will not significantly impact the strength of their results, when in reality, the opposite occurs. Rather than increasing the power to detect the effects of stress, the effect they make more robust is, in fact, the effect of the data treatment strategy itself.

It can be argued that the increased false positive rate we found in our simulation only applies to 41% of the simulated experiments (what we called viable experiments), and that there is a much lower risk of finding a false positive result (20% or 29% for t-test and ANOVA, respectively) when the non-viable experiments are factored in. While this may be the case, we would like to sustain that employing these data treatment strategies based on loosely defined criteria still pose a risk. To support our claim we prompt the reader to consider what “non-viable experiments” might mean in the real world. Non-viable experiments were those in which less than three simulated rats presented with a preference below the threshold that defines susceptibility (in our case, 65%). Statistical comparisons after data treatment would not be possible in such a situation. How would a researcher respond to a situation like this if they were encountering this in their stress model? We also need to consider why it is that we have not identified any studies like this in our literature investigations. A few possible scenarios come to mind. Ideally, non-viable experiments could not happen in the laboratory because stress always has a deleterious effect on sweet preference. The effect is always of a magnitude such that it decreases the outcome below the desired threshold. This is an unlikely scenario from our point of view. In our systematic review we analyzed studies reporting little to no change in sweet preference after stress^6^. In some studies the preference for sucrose even increased^51,52^ Even in this ideal scenario, we risk overestimating the effect of stress as explained above. The alternative scenarios far even worse, however. We could, for example, imagine that non-viable experiments *do* happen, but results are simply not published. This is sometimes referred as the “file drawer problem”^53^ and a possible explanation to the significant small study effect in the form of funnel plot asymmetry^54^ we found in our systematic review^6^. In this imagined situation, the true false positive rate for using this type of data treatment is unknowable. A practical response might be to increase the sample size – add more rats, addressing the problem of having too few susceptible individuals – in order to turn the experiment “viable.” Adding subjects like this to a study is problematic and known to increase false positive rates^55^. In this specific scenario it is doubly problematic since, as we explained before, the false positive rates of the data treatment strategies increase, rather than decrease, with larger experiments. Other practical approaches are similarly problematic. An insufficient number of animals with a preference below a predefined threshold might prompt a *post hoc* adjustment of the cut-off to fit the data, thus salvaging the experiment. All scenarios, to some extent, contribute to a biased representation of the effect of stress on sweet consumption, increasing the risk of reaching misleading conclusions in one’s experiments. With this in mind, researchers need to be aware of the unintended consequences of their data treatment strategies.

## Conclusion

By making use of the sucrose preference test, some researchers discriminate between individuals susceptible and resilient to stress by defining a threshold based on their sucrose consumption. This threshold varies from study to study, which may have major implications with respect to the conclusions drawn from the experiments. In our descriptive study, we found that these thresholds were not only varied, but mostly not evidence-based. Most authors either refrained from providing a justification for their methods or relied exclusively on previous publications. When tracing these citations, we converged on a pioneering article that ultimately could not be regarded as supporting the methods used in studies that had referenced it. In fact, the study had not even been carried out in rats. Indeed, the idea of a universal cut-off for anhedonia seems counterintuitive given rats’ phenotypic heterogeneity in sweet consumption, the methodological flexibility of the model and the unproven assumption that the test is a reliable measure of general anhedonia. What is more, researchers might unintendedly introduce statistical bias when deciding either to remove or split data based on this predefined threshold. Through a simulation study, we provided concrete evidence that these data treatment strategies confound the results of experiments by creating artificial differences also where there are none. This was true also when we used large sample sizes. Increasing the sample size paradoxically reinforced the bias created by the data treatment strategies. We would like to caution researchers against implementing a specific cut-off in the sucrose preference test to define anhedonia based solely on previous literature. If a threshold is used, researchers need to be mindful of the potential unintended consequences of their data treatment strategy. We also call for more transparent reporting of sucrose preference test methods. The reasoning behind methodological decisions and changes from previous studies need to be clearly explained.

