## Supplementary material for "Rethinking data treatment: The sucrose preference threshold for anhedonia in stress-induced rat models of depression"

### Supplementary material

Jenny P. Berrio[1], Otto Kalliokoski[1]

#### Methods

##### Registration and open access data

Access to the code, additional files and data sets can be found here: https://osf.io/egk8s/
Throughout the text you will find references to specific files found in this repository.

##### Figure 1 copyright attribution

Water Bottle by DT. Creative Lab, sugar by Laymik, Mouse by Cynthia Tran Vo, Heart by BomSymbols, drop by Baboon designs and Arrow by kiddo from Noun Project (CCBY3.0).

Arrow by kiddo from Noun Project (CCBY3.0) --> Supplementary Figure 3, 4, 7, 8, 10

##### Deviations from protocol

Initially, we had planned to use three different datasets (derived from two original studies) to calculate the sweet and water consumptions to be used in our simulations. However, in an effort to obtain values that more closely reflect the variability of real-world data, we decided to use our systematic review dataset instead, as described below.

##### Back-tracing study

Access to the data: *(OSF: "Backtracing.xlsx")*

##### Simulation study

The following R packages were used in the simulation study:

#### Iterations

In order to select the number of simulated experiments that would provide the most accurate estimate of the false positive rate, we performed a series of simulations where either 10,000, 100,000 or 1,000,000 experiments were analyzed. A million experiments were needed for an accurate estimate of the false positive rate (Supplementary table 1) and was chosen for the study.

**Supplementary table 1**. Series of iterations of 10,000, 100,000 or 1,000,000 simulated experiments.

| # Experiments | Iteration | t-test Normal experiments | t-test Data exclusions | ANOVA Data split |
|---|---|---|---|---|
| 10,000 | 1 | 0.049 | 0.208 | 0.118 |
| 10,000 | 2 | 0.050 | 0.215 | 0.125 |
| 10,000 | 3 | 0.047 | 0.212 | 0.124 |
| 10,000 | 4 | 0.048 | 0.210 | 0.119 |
| 10,000 | 5 | 0.046 | 0.215 | 0.122 |
| 10,000 | 6 | 0.046 | 0.220 | 0.121 |
| 10,000 | 7 | 0.052 | 0.213 | 0.119 |
| 10,000 | 8 | 0.050 | 0.218 | 0.121 |
| 10,000 | 9 | 0.048 | 0.219 | 0.124 |
| 10,000 | 10 | 0.047 | 0.210 | 0.117 |
| | **Mean** | **0.048** | **0.214** | **0.121** |
| | **Standard deviation** | **0.002** | **0.004** | **0.003** |
| 100,000 | 1 | 0.049 | 0.213 | 0.122 |
| 100,000 | 2 | 0.047 | 0.215 | 0.125 |
| 100,000 | 3 | 0.049 | 0.213 | 0.123 |
| 100,000 | 4 | 0.048 | 0.215 | 0.123 |
| 100,000 | 5 | 0.048 | 0.213 | 0.123 |
| 100,000 | 6 | 0.047 | 0.215 | 0.124 |

| | | | | |
|---|---|---|---|---|
| 100,000 | 7 | 0.047 | 0.214 | 0.122 |
| 100,000 | 8 | 0.049 | 0.212 | 0.122 |
| 100,000 | 9 | 0.047 | 0.213 | 0.122 |
| 100,000 | 10 | 0.048 | 0.213 | 0.123 |
| | **Mean** | **0.048** | **0.214** | **0.123** |
| | **Standard deviation** | **0.001** | **0.001** | **0.001** |
| 1,000,000 | 1 | 0.0477 | 0.2126 | |
| 1,000,000 | 2 | 0.0480 | 0.2133 | |
| 1,000,000 | 3 | 0.0476 | 0.2135 | |
| 1,000,000 | 4 | 0.0479 | 0.2132 | |
| 1,000,000 | 5 | 0.0478 | 0.2131 | |
| | **Mean** | **0.0478** | **0.2132** | |
| | **Standard deviation** | **0.0002** | **0.0003** | |

#### Calculation of consumptions and preference – simulation study

Access to the data: *(OSF: "Database_main.xlsx")*

To obtain average values for sucrose and water consumption of unstressed rats in the sucrose preference test, we used our systematic review dataset (REF!). First, we extracted preference values for 1-2% sucrose in 1-hour tests for unstressed rats and synthesized them using random effects models (package "dmetar" (Harrer et al., 2019)). Forty-eight mean values were combined and a pooled value of 82.1% (95% CI: 80.01%, 84.15%) was obtained (Supplementary figure 1). In order to obtain an averaged value of sweet intake with its measure of dispersion, 14 studies that reported intake (g or mL) of 1-2% sucrose in 1-hour tests were combined according to the Cochrane handbook (Higgins et al., 2022). Based on these studies, an intake of 12.22 ± 7.49 mL sucrose solution was obtained. Using the obtained values for preference and sucrose intake allowed us to solve for an estimate of the water intake. Water intake was assumed to have the same coefficient of variation as the sucrose intake. Based on this, an averaged water intake of 2.66 ± 1.63 mL was obtained. An averaged leakage per bottle of 1 to 3 droplets (water droplet have an approximate volume of 0.05 mL) was assumed for the model.

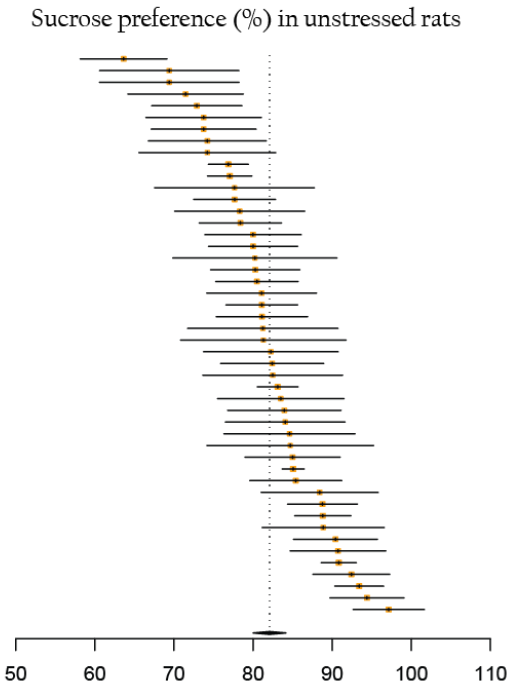


**Supplementary figure 1.** Simplified forest plot of the preference for 1-2% sucrose in 1-hour sucrose preference tests of 48 unstressed cohorts of rats. Each line represents one cohort, with its mean (yellow square) and its standard error (length of the line). High variation in the results was observed (between-study heterogeneity: $I^2$ = 87.3% %; 95% CI: 84%; 89.9%). The dotted line represents the pooled preference and the width of the black diamond its 95% confidence interval.

#### Normal experiments – increased sample size

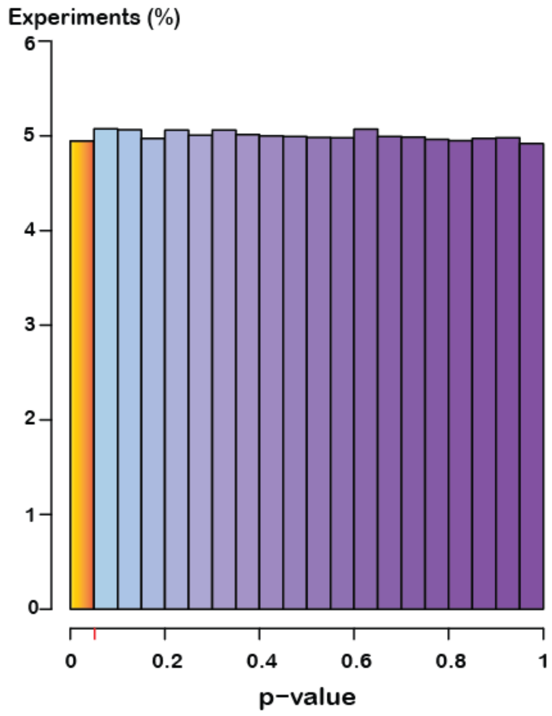


**Supplementary figure 2. False positive rates normal experiments with increased sample size. A.** Normal experiments in which unmodified cohorts of 100 rats with simulated sucrose preference values were compared to each other using Student's t-test. It is a well-established fact that the t-test works well also with data that deviate from a normal distribution (Box, 1953) and it is probably still the best choice for our analyses despite the slightly conservative false positive rate estimates. False positive rate: 0.049. Red marker on the x-axis/yellow bar represent a p value of 0.05.

#### Shiny app- Sweet preference test simulator

Access to the code: https://osf.io/g9bxs/

The following R packages were used in the developing the app:

In the following section, you will find detailed instructions on how to use the app. For further inquiries, write us at.

In the apps initial page (Supplementary figure 3), you will find a brief introduction explaining the rationale and purpose of the app. This background information can be concealed or shown at will. To the far right, two tabs will provide you with extra details. A small help section will give you simple instructions. An “About” section will provide you with contact details and with links to the OSF and GitHub repositories hosting the application code. To the far left, you will find the main part of the app – The sweet preference test simulator - where you will be able to either use the values assumed for our model or type in your own data.

The simulator is composed of two main sections (supplementary figure 4). In the section on the left (“Consumptions”) you will be able to modify the average sweet and water consumptions, along with their respective standard deviations. There is also space to include the average leak of the bottles in your lab.

In the section to the right (“Experiment”), you will be able to select the parameters of the experiments you want to simulate. That is, the number of animals in each of the cohorts, the preference threshold you want to employ to distinguish between susceptible and resilient animals, and the number of experiments to simulate (on a range between 100 and 1,000). Be aware that the cohort named “stressed” will be the one modified according to the selected preference threshold. Per default, the consumptions and experiment parameters assumed in our study are selected.

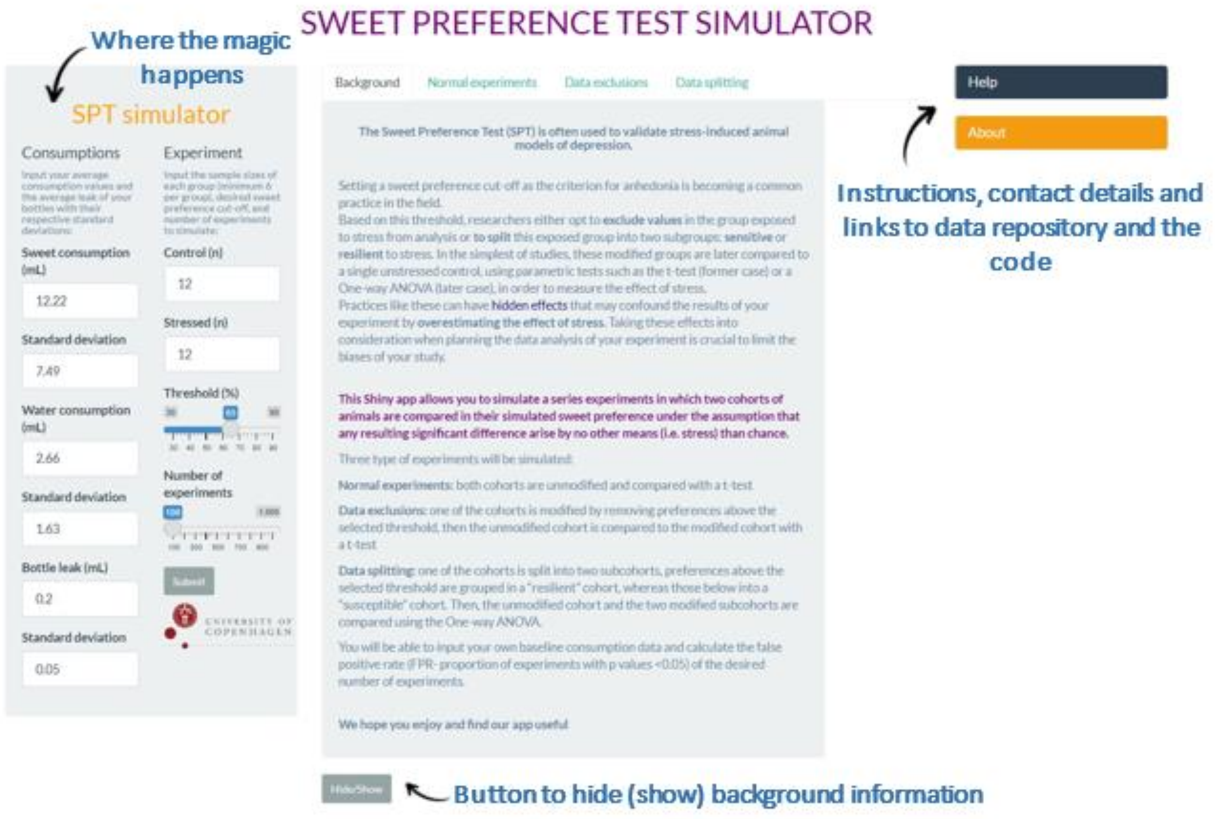


**Supplementary figure 3. Front page of the Shiny app.** This is an image of the elements on the front page of the app. Pointed out with arrows are the main sections where the background information, instructions, app code, and contact details can be found. To the left side you will find the sweet preference simulator.

As observed in supplementary table 1, 1,000 experiments can give a good approximation of the false positive rate; however, their computation requires more time. This is particularly the case for graphs where the false positive rate is calculated as the sample size or the threshold changes. In these instances, we ask you to be patient. A bar indicating the progress will be shown to you in the bottom right.

Once you are satisfied with the parameters of the simulations, go to the tab of your interest and click **Submit.** Each tab corresponds to one of the three different scenarios simulated in our study. The graphs that do not require any additional information will be shown automatically with the parameters selected once you click submit.

The "Normal experiments" tab (Supplementary figure 5) will show you a histogram of the p values of the t-test comparing two unstressed cohorts of rats with their simulated preferences. The "Data exclusions" tab has three sub-tabs. The p-values distribution sub-tab (Supplementary figure 6) will plot the p values of the t-test comparing experiments where one of the cohorts (“stressed“) is modified by removing data points above the selected preference threshold. The false positive rate and the proportion of experiments where the comparison was not possible is shown below the histogram. Be aware that the false positive rate calculated here is that of the viable experiments, and not of the total number of experiments simulated.

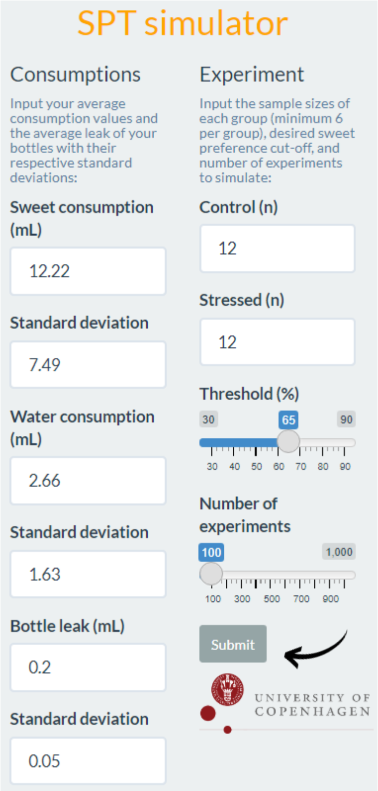


**Supplementary figure 4. Sweet preference test simulator.** The simulator is composed of two sections. In the section “Consumptions”, the average amount of sweet and water intake can be adjusted to any desired values. It is also possible to adjust the average leakage of the bottles (this will often be proportional to the duration of the test). Be aware that this parameter will be applied to both the water and sweet consumptions (independently) when the sweet preferences are simulated. In the section “Experiment”, you can select the number of animals per cohort, preference threshold, and number of experiments to simulate. Click **Submit** to start the calculations.

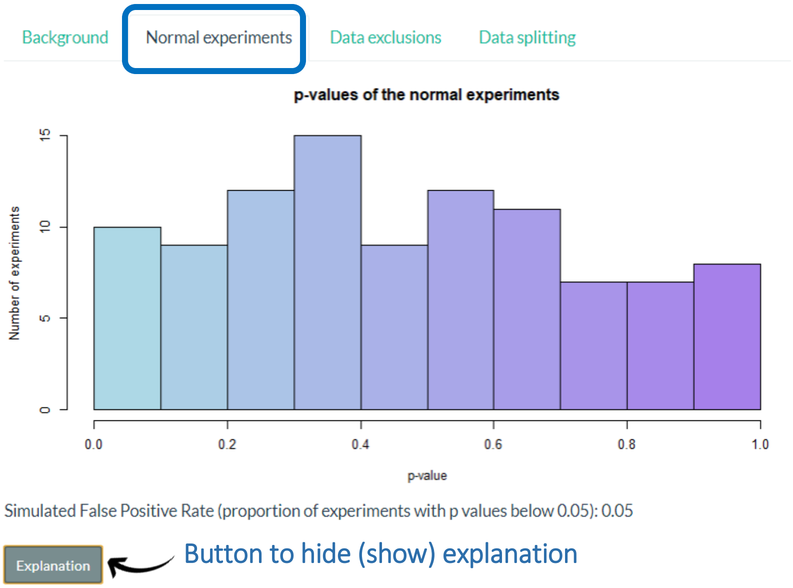


**Supplementary figure 5. Normal experiments tab.** A histogram of the p values of the t-tests is shown along with the false positive rate (proportion of experiments with a p value below 0.05). By clicking on the button “Explanation”, you can show or hide additional information detailing the parameters of the experiment.

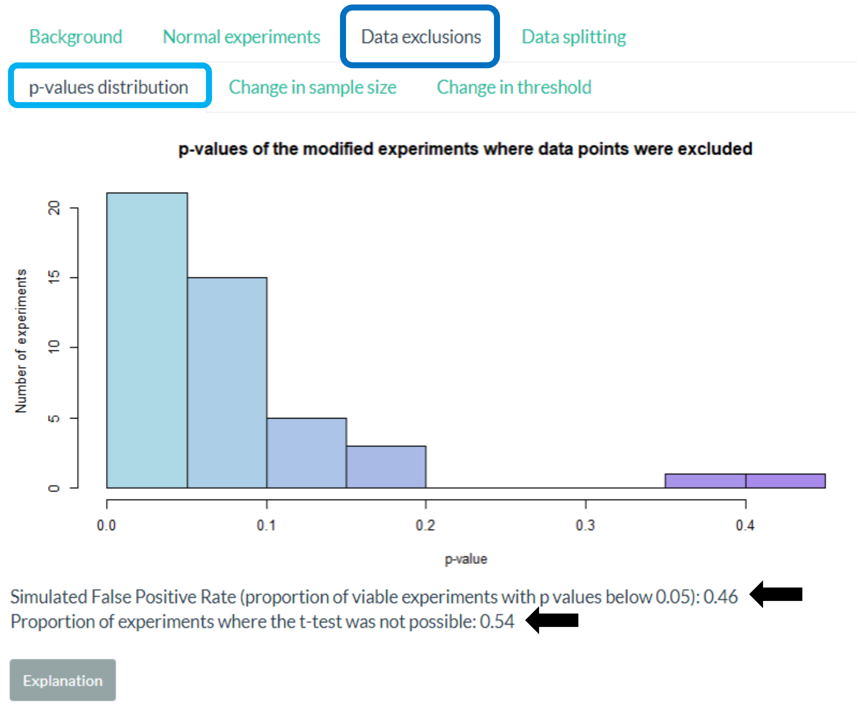


**Supplementary figure 6. Data exclusions – p-values distribution tab.** A histogram of the p values of the t-tests of viable experiments is shown along with the false positive rate and the proportion of experiments where the comparison was not possible because less than three values were left in the "stressed" cohort (arrows). By clicking on the button "Explanation", you can show or hide additional information detailing the parameters of the experiment.

The Change in sample size and Change in threshold sub-tabs will plot the false positive rates as a function of sample sizes and the preference thresholds. In each of these sub-tabs you will find an additional box where you will be able to select the ranges of values to plot (Supplementary figure 7). Once you have set your parameters, click **Go.** Contrary to the p values distribution tab, the calculations of the false positive rates in these sections are based on the total number of experiments you chose to simulate. The code for these plots was made so non-viable experiments are skipped until the desired number of viable experiments to simulate is reached. This is the reason why these two plots take the longest to appear.

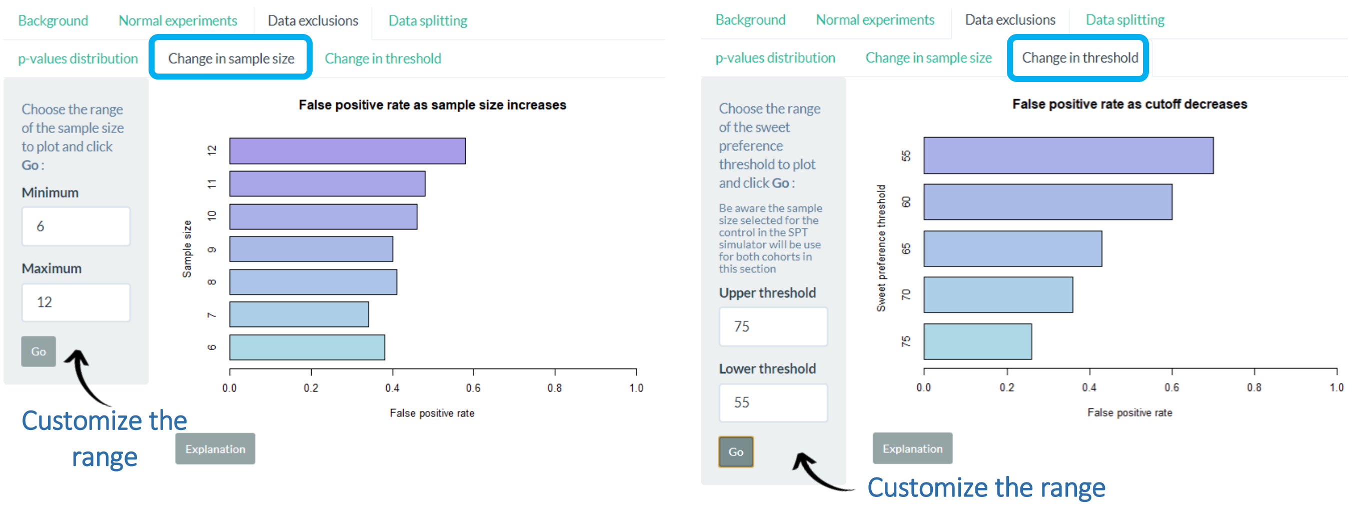

**Supplementary figure 7. Data exclusions – change in sample size and threshold sub-tabs.** The false positive rate of the simulated experiments is plotted against the initial sample size of the cohorts or the preference threshold. An additional box to the left allows you to customize the range to plot. Click **Go** to render the plots. By clicking on the button "Explanation", you can show or hide additional information detailing the parameters of the experiment.

Likewise, the "Data splitting" tab has four supplementary tabs. The first one ("ANOVA") provides an example experiment where one of the cohorts ("stressed") is split in two subgroups. Subjects with preference values above the selected threshold form a "resilient" subgroup, while subjects with values below formed the "susceptible" subgroup. These two subgroups and a control (unmodified) cohort are compared using a one-way ANOVA, followed by Tukey´s (HSD) post hoc test, if significant (Supplementary figure 8). Below the boxplot, you will see the results of the statistical test. In cases where the comparison is not possible, the message "One of the groups to compare has less than three data points. The statistical analysis is not possible" will be shown instead. To generate a new example, simply click **Submit**.

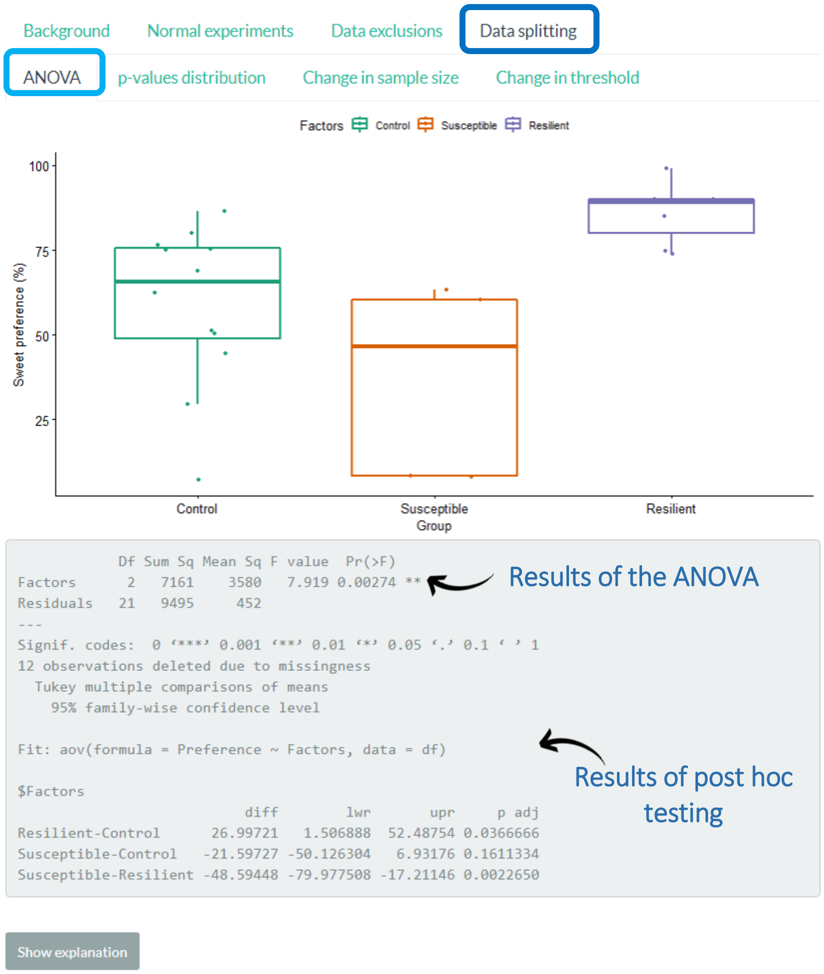


**Supplementary figure 8. Data splitting – ANOVA tab.** Boxplot of an example experiment where the stressed group is split into susceptible and resilient subjects. The results of the ANOVA/*Post hoc* testing are displayed below the graph. By clicking on the button "Explanation", you can show or hide additional information detailing the parameters of the experiment. Click **Submit** if you want to generate a new example.

The following three sub tabs are similar to those previously described for the Data exclusion tab. The only difference is that two plots, instead of one, are rendered in each sub-tab. The first plot shows the results of the ANOVA, while the second plot shows the *post hoc* test. In the p-values distribution sub-tab, you will be able to see either the distributions of the p values of the ANOVA test or that of Tukey´s *post hoc* test comparing the controls with the susceptible group. The false positive rate and the proportion of non-viable experiments is reported for the ANOVA, while only the false positive rate is reported for the *post hoc* tests. In these tabs, the calculated false positive rate is that of the proportion of viable experiments, and not of the total number of experiments simulated. In the change in sample size and threshold tabs, the two different plots can be selectively shown using a drop down list (Supplementary figure 10). Similar to those in “Data exclusions”, the calculations of the false positive rates are based on the total number of experiments simulated.

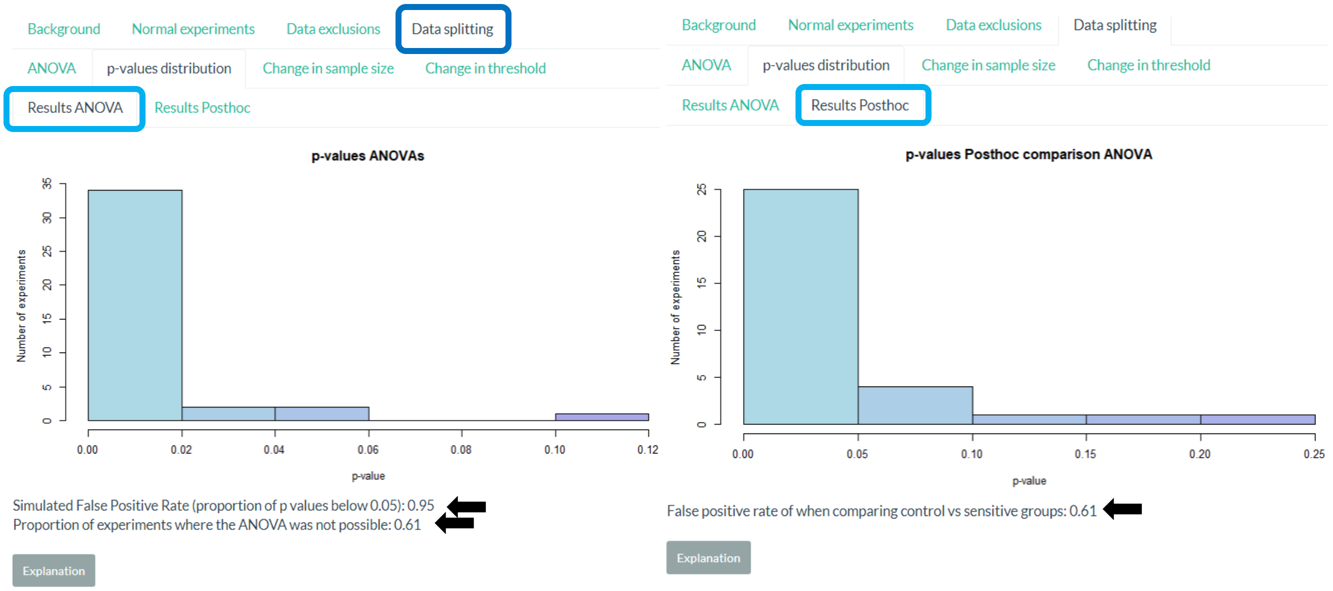


**Supplementary figure 9. Data splitting – p-values distribution tab**. Two different histograms are plotted here. In the “Results ANOVA” tab, the p values of the ANOVA test of viable experiments is shown along with the false positive rate and the proportion of experiments where the comparison was not possible (arrows). In the “Results Posthoc” tab, the p values are that of Tukey’s *post hoc* tests comparing the “control” and the “susceptible” groups. The false positive rate is reported below the histogram (arrow). By clicking on the button “Explanation”, you can show or hide additional information detailing the parameters of the experiment.

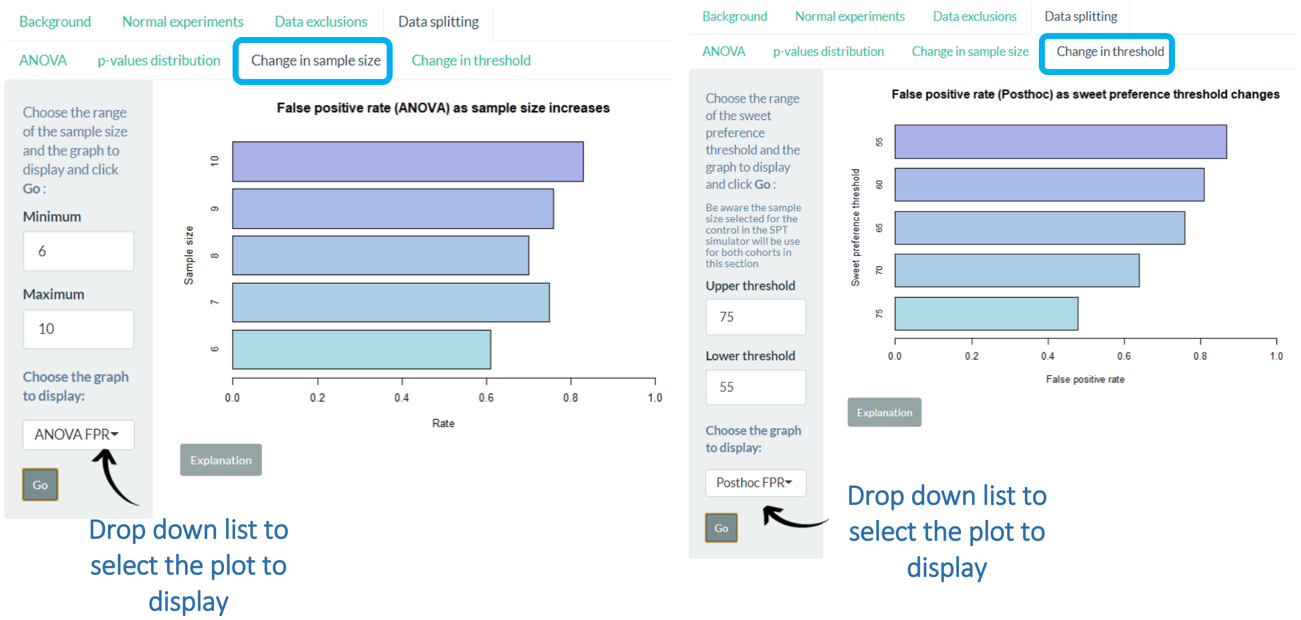


**Supplementary figure 10. Data splitting – change in sample size and threshold tabs.** The false positive rate of viable experiments is plotted against the initial sample sizes of the cohorts or the preference thresholds. An additional box to the left allows you to customize the range of values to plot. A dropdown list allows you to select between two different plots: one displaying the false positive rates of the ANOVAs ("ANOVA FPR") or those of the *post hoc* tests ("Posthoc FPR"). Click **Go** to render the plots. By clicking the button "Explanation", you can show or hide additional information detailing the parameters of the experiment.
